# GABA decrease is associated with degraded neural specificity in the visual cortex of glaucoma patients

**DOI:** 10.1101/2022.10.09.508561

**Authors:** Ji Won Bang, Carlos Parra, Kevin Yu, Gadi Wollstein, Joel S. Schuman, Kevin C. Chan

## Abstract

Glaucoma is an age-related neurodegenerative disease of the visual system, affecting both the eye and the brain. Yet its underlying metabolic mechanisms and neurobehavioral relevance remain largely unclear. Here, using proton magnetic resonance spectroscopy and functional magnetic resonance imaging, we investigated the GABAergic and glutamatergic systems in the visual cortex of glaucoma patients, as well as neural specificity, which is shaped by GABA and glutamate signals and underlies efficient sensory and cognitive functions. Our study showed that among the older adults, both GABA and glutamate levels decrease with increasing glaucoma severity regardless of age. Further, the reduction of GABA but not glutamate predicted the neural specificity. This association was independent of the impairments on the retina structure and age. Our results suggest that glaucoma-specific decline of GABA undermines neural specificity in the visual cortex and that targeting GABA could improve the neural specificity in glaucoma.

## Introduction

Glaucoma is an age-related neurodegenerative disease characterized by the gradual loss of retinal ganglion cells. Such damage disrupts the transmission of the retinal signals to the brain and eventually leads to irreversible blindness. With the increasing aging population, it is estimated that 111.8 million people worldwide will be affected by glaucoma in 2040, suggesting that glaucoma is a major public health problem^1^. Nevertheless, the exact mechanisms underlying glaucoma remain unclear. Current clinical treatments focus on intraocular pressure (IOP) reduction with laser, medication, or surgery. However, IOP is a major risk factor but not the cause of the disease. Glaucoma progression in certain patients cannot be attenuated even with controlled IOP, indicating that glaucoma cannot be explained by IOP elevation alone. To reduce the prevalence of this irreversible but preventable disease, we need a better understanding of the mechanisms of glaucoma that will lead to improved treatment strategies beyond IOP control.

An increasing amount of studies has pointed out that glaucomatous degeneration occurs along the visual pathway including the optic nerves^2,3^, optic chiasm^3^, lateral geniculate nucleus^3–9^, optic radiation^2,10^, and visual cortex^3,6,7,9,11–13^. Further, recent studies have suggested that glaucoma may share some pathogenic mechanisms with Alzheimer’s disease, which is characterized by the accumulation of amyloid β and tau^14,15^. Critically, glaucoma patients were shown to present abnormal tau proteins in the retina^16^ and vitreous fluid^17^. Likewise, in animal models of glaucoma, amyloid β and tau were found in the retina^18^, lateral geniculate nucleus, and even in the primary visual cortex^19^, although the amount of deposits was relatively lower in the visual cortex than in the anterior visual pathway. It suggests that the disease may spread trans-synaptically from the anterior to posterior visual pathways^19^.

Accumulation of amyloid β and tau in Alzheimer’s disease was found to impair the glutamatergic and GABAergic systems, which are the main excitatory and inhibitory systems in the brain. Exposure to amyloid β can increase extracellular glutamate levels by preventing glutamate uptake^20,21^ or potentiating glutamate release^22,23^. Elevated amyloid β and tau can also impair synaptic inhibition via downregulation of GABA_A_ receptor^24^ and induce significant loss of GABAergic neurons^25–27^ and GABA-synthetic enzyme^25^. Of note, loss of GABAergic interneurons co-localizes with tau markers^25^. If glaucoma patients present amyloid β and tau accumulation in the visual system of the brain, the glutamatergic and/or GABAergic signals are likely to be impaired similar to individuals with Alzheimer’s disease.

In particular, the GABAergic signals play a critical role in shaping the response patterns of neuronal populations^28^. Recent magnetic resonance spectroscopy (MRS) studies revealed a tight relationship between GABAergic signals and neural specificity at the level of neuronal population. As the cortical GABA levels decrease, neural activity patterns associated with different categories become similar, thus more confusable^29,30^. Increased confusability of neural activity patterns then could potentially degrade behavioral performance^31^. Indeed, loss of neural specificity was observed to be associated with declines in memory performance^32^ and fluid processing abilities such as executive function and speed of processing^33^. Relatedly, glaucoma patients were found to present impairments on some of the perceptual and cognitive functions where GABAergic signals are thought to be involved. These include abnormalities in visual crowding effect^34^, binocular rivalry^35–38^, visual categorization^39,40^, visual attention^41^, and cognitive capacity^42^. Further, glaucoma is thought to accelerate the speed of aging process^43^, which involves gradual reduction of cortical GABA levels^44–49^. It suggests that glaucomatous degeneration may involve the reduction of GABA to a greater extent than healthy aging and that this reduction of GABA may affect the neural specificity. Recent hypothesis-independent pathway analysis also implicates GABA and acetyl-CoA metabolism as the only pathway that is significantly associated with both primary open-angle glaucoma and normal-pressure glaucoma^50^. Nevertheless, other than the limited studies on GABAergic involvements in the retina of experimental glaucoma models^51,52^, no studies have directly examined glaucoma-specific changes in the GABAergic signals and their relationship with neural specificity in the visual cortex of glaucoma patients.

Therefore, in the current study, we investigated whether the GABA and glutamate levels in the visual cortex are affected by glaucoma and whether these neurochemical changes are associated with the neural specificity independent of the impairments of the retina structure and age. To address these questions, we recruited glaucoma patients and age-matched healthy subjects and conducted functional magnetic resonance imaging (fMRI), MRS of the brain, as well as clinical ophthalmic tests including Humphrey visual field perimetry and optical coherence tomography (OCT) of the retina and optic nerve. Using principal component analyses (PCA) and regression modeling, we observed that among our older adult subjects, GABAergic and glutamatergic signals in the visual cortex gradually reduced with increasing severity of glaucoma regardless of age. Further, we observed that the neural specificity in the visual cortex was tightly related to the glaucoma-specific reduction of GABAergic signals, but not that of glutamatergic signals. This association between GABA and neural specificity was observed independent of age or impairments on the retina. These findings indicate the importance of the GABAergic action in the visual cortex and its implications in sensory encoding in glaucoma.

## Results

Forty-one glaucoma patients and twenty-three age-matched healthy subjects underwent clinical ophthalmic exams and magnetic resonance imaging. Their demographic and clinical characteristics can be found in **Table 1**. Specifically, we obtained each individual’s structural and functional images of the whole brain, neurochemical profiles of the visual cortex (**Fig. 1**) as well as clinical ophthalmic measures including peripapillary retinal nerve fiber layer (pRNFL) thickness, macular ganglion cell-inner plexiform layer (mGCIPL) thickness, optic nerve head cup-to-disc (C/D) ratio, and neuroretinal rim (NRR) area from OCT and the visual field mean deviation (MD) from Humphrey standard automated perimetry.

**Fig. 1:**
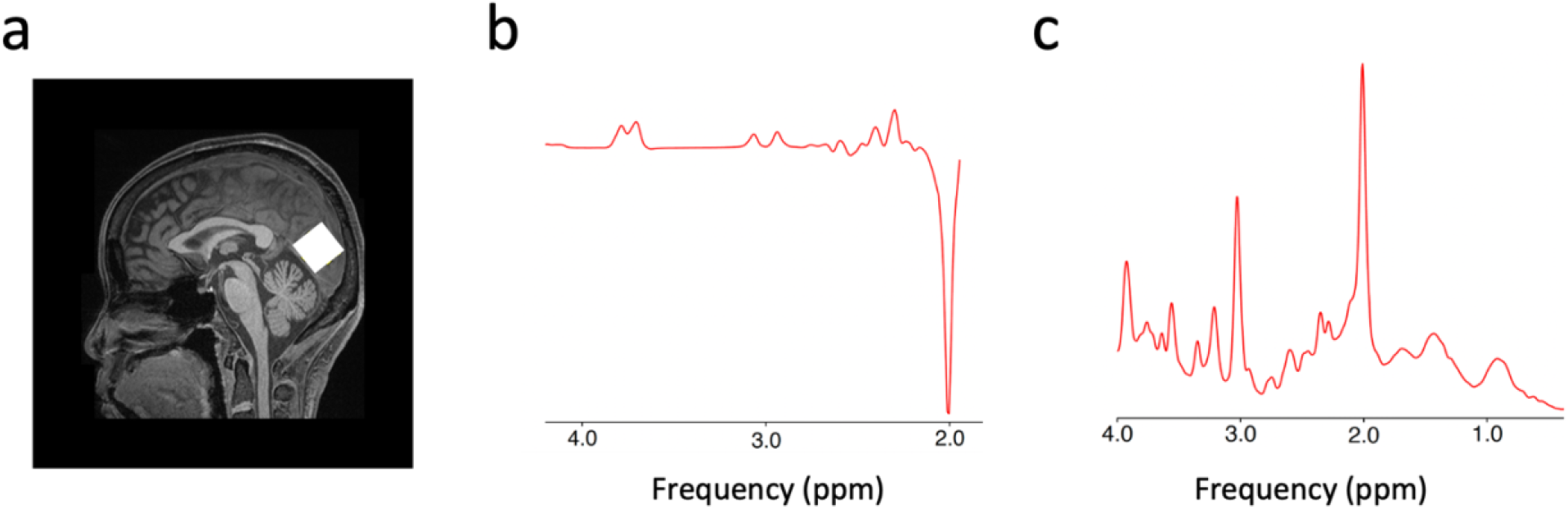
Sample voxel location for proton MRS and representative spectra for GABA and glutamate. **a,** A 2.2 x 2.2 x 2.2 cm^3^ voxel (white box) was manually positioned along the calcarine sulci on the occipital lobe. **b,** A sample spectrum from the MEGA-PRESS sequence for GABA. The GABA peak is at 2.8-3.2 ppm. **c,** A sample spectrum from the PRESS sequence for glutamate. The glutamate peak is at 2.2-2.4 ppm.

**Table 1.** Demographic and clinical characteristics of the glaucoma and healthy subjects.

| | Healthy control<br>(mean ± S.E.M.) | Early glaucoma<br>(mean ± S.E.M.) | Advanced glaucoma<br>(mean ± S.E.M.) | Group effect<br>(one-way ANOVA or Chi-squared test ) |
| --- | --- | --- | --- | --- |
| Age (year) | 64.78 ± 1.63 | 65.33 ± 2.25 | 66.19 ± 1.47 | F(2,61)=0.198, $P=0.821$ |
| Sex | 11 M, 12 F | 4 M, 11 F | 13 M, 13 F | $\chi^2(2)=2.347$, $P=0.309$ |
| Average pRNFL thickness ($\mu\text{m}$) OD | 88.68 ± 2.11 | 73.79 ± 3.49 | 62.04 ± 1.95 | F(2,56)=35.203, $P<0.001$ |
| Average pRNFL thickness ($\mu\text{m}$) OS | 91.05 ± 1.94 | 77.79 ± 2.95 | 64.62 ± 2.07 | F(2,56)=38.227, $P<0.001$ |
| mGCIPL thickness ($\mu\text{m}$) OD | 79.25 ± 1.52 | 68.62 ± 3.67 | 59.52 ± 2.10 | F(2,49)=18.495, $P<0.001$ |
| mGCIPL thickness ($\mu\text{m}$) OS | 78.56 ± 2.13 | 72.15 ± 2.47 | 58.57 ± 1.50 | F(2,49)=31.296, $P<0.001$ |
| NRR area ($\text{mm}^2$) OD | 1.24 ± 0.04 | 0.97 ± 0.13 | 0.64 ± 0.04 | F(2,56)=23.242, $P<0.001$ |
| NRR area ($\text{mm}^2$) OS | 1.34 ± 0.04 | 0.95 ± 0.11 | 0.66 ± 0.04 | F(2,56)=36.588, $P<0.001$ |
| Optic nerve head C/D OD | 0.44 ± 0.04 | 0.66 ± 0.06 | 0.75 ± 0.02 | F(2,56)=21.698, $P<0.001$ |
| Optic nerve head C/D OS | $0.42 \pm 0.04$ | $0.65 \pm 0.05$ | $0.75 \pm 0.03$ | $F(2,56)=26.975, P<0.001$ |
| Visual field mean deviation (dB) OD | $-1.72 \pm 0.63$ | $-2.00 \pm 0.58$ | $-17.36 \pm 1.65$ | $F(2,55)=48.930, P<0.001$ |
| Visual field mean deviation (dB) OS | $-1.50 \pm 0.43$ | $-2.44 \pm 0.63$ | $-14.33 \pm 1.67$ | $F(2,55)=31.182, P<0.001$ |

Abbreviations: pRNFL: peripapillary retinal nerve fiber layer; mGCIPL: macular ganglion cell-inner plexiform layer; NRR: neuroretinal rim; C/D: cup-to-disc; OD: oculus dexter (right eye); OS: oculus sinister (left eye).

To examine whether the GABAergic system in the visual cortex alters with disease severity, we first compared the GABA levels across healthy controls, early glaucoma subjects (average MD between eyes better than −6.0dB) and advanced glaucoma subjects (average MD between eyes worse than −6.0dB). A one-way ANCOVA while regressing out the effect of age revealed a significant main effect of group (F(2,54)=7.162, *P*=0.002, partial η^2^=0.210; **Fig. 2a**). Post-hoc tests showed that the levels of GABA are significantly reduced in advanced glaucoma patients compared to healthy controls (early glaucoma vs. healthy controls, Bonferroni-corrected *P*=0.203, 95% CI=-0.070 – 0.010, advanced glaucoma vs. healthy controls, Bonferroni-corrected *P*=0.001, 95% CI=-0.086 – −0.018, early glaucoma vs. advanced glaucoma, Bonferroni-corrected *P*=0.534, 95% CI=-0.018 – 0.061).

**Fig. 2:**
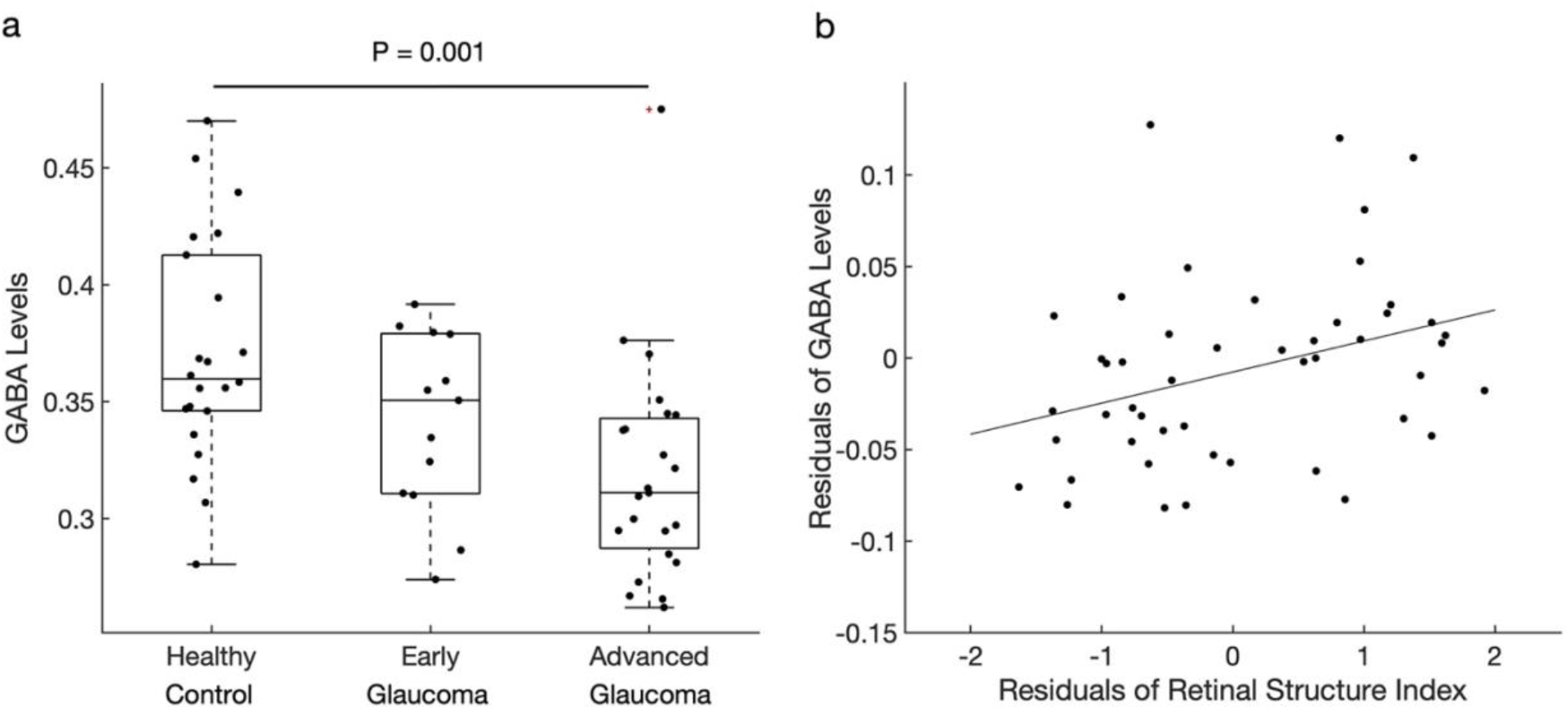
GABA levels in the visual cortex. **a,** The GABA levels (normalized to total creatine levels) were significantly reduced in advanced glaucoma patients compared to healthy controls (Bonferroni-corrected *P*=0.001). The distributions are represented using box and whisker plots and the outlier is plotted as a red plus sign. In descending order, the lines in the plots represent: maximum, third quartile, median, first quartile, and minimum. Individual data points are overlaid on the box plots. **b,** Individual differences in the retinal structure index were significantly associated with individual differences in the GABA levels after controlling for age (r=0.339, *P*=0.020).

Next, we also tested whether the impairments on the retina structure, which is a fine-grained, continuous measure of disease severity in the eye are associated with the reduction of GABA levels in the visual cortex. For this, we obtained the marker for retinal structural damage by extracting a common component from each individual’s pRNFL thickness, mGCIPL thickness, C/D ratio, and NRR area of left and right eyes using PCA. The PCA yielded only one component that had an eigenvalue greater than one, explaining 73.42% of variances of all clinical OCT measures. Using this component, termed as retinal structure index, we ran a linear regression analysis to estimate the GABA levels. The results showed that the retina structure significantly predicted the GABA levels while controlling for age (T(45)=2.414, *P*=0.020, β=0.337, partial correlation=0.339, R^2^ change=0.113; **Fig. 2b**). It indicates that those who have greater impairments on the retina structure (i.e., lower value of the retinal structure index) present lower amounts of GABA in the visual cortex independent of their ages. The effect of age, however, failed to predict the amount of GABA after controlling for the retinal structure index (T(45)=0.596, *P*=0.554, β=0.083, partial correlation=0.088, R^2^ change=0.007).

Next, we examined whether the glutamate levels exhibit similar changes as GABA levels in glaucoma. A one-way ANCOVA with a factor of group controlling for age revealed a significant main effect of group (F(2,56)=5.079, *P*=0.009, partial η^2^=0.154; **Fig. 3a**). The levels of glutamate were significantly lower in advanced glaucoma patients compared to healthy controls and early glaucoma patients (early glaucoma vs. healthy controls, Bonferroni-corrected *P*=1.000, 95% CI=-0.120 – 0.139, advanced glaucoma vs. healthy controls, Bonferroni-corrected *P*=0.026, 95% CI=-0.242 - −0.012, early glaucoma vs. advanced glaucoma, Bonferroni-corrected *P*=0.030, 95% CI=0.010 - 0.263). Further, the linear regression analysis demonstrated that the retinal structure index significantly predicted the levels of glutamate while the effect of age was controlled (T(46)=2.654, *P*=0.011, β=0.358, partial correlation=0.364, R^2^ change=0.128; **Fig. 3b**). It suggests that older adults with greater impairments on the retina structure present lower amounts of glutamate in the visual cortex independent of their ages. Nevertheless, age failed to account for the levels of glutamate when the effect of the retina structure was controlled (T(46)=-1.525, *P*=0.134, β=-0.206, partial correlation=-0.219, R^2^ change=0.042).

**Fig. 3:**
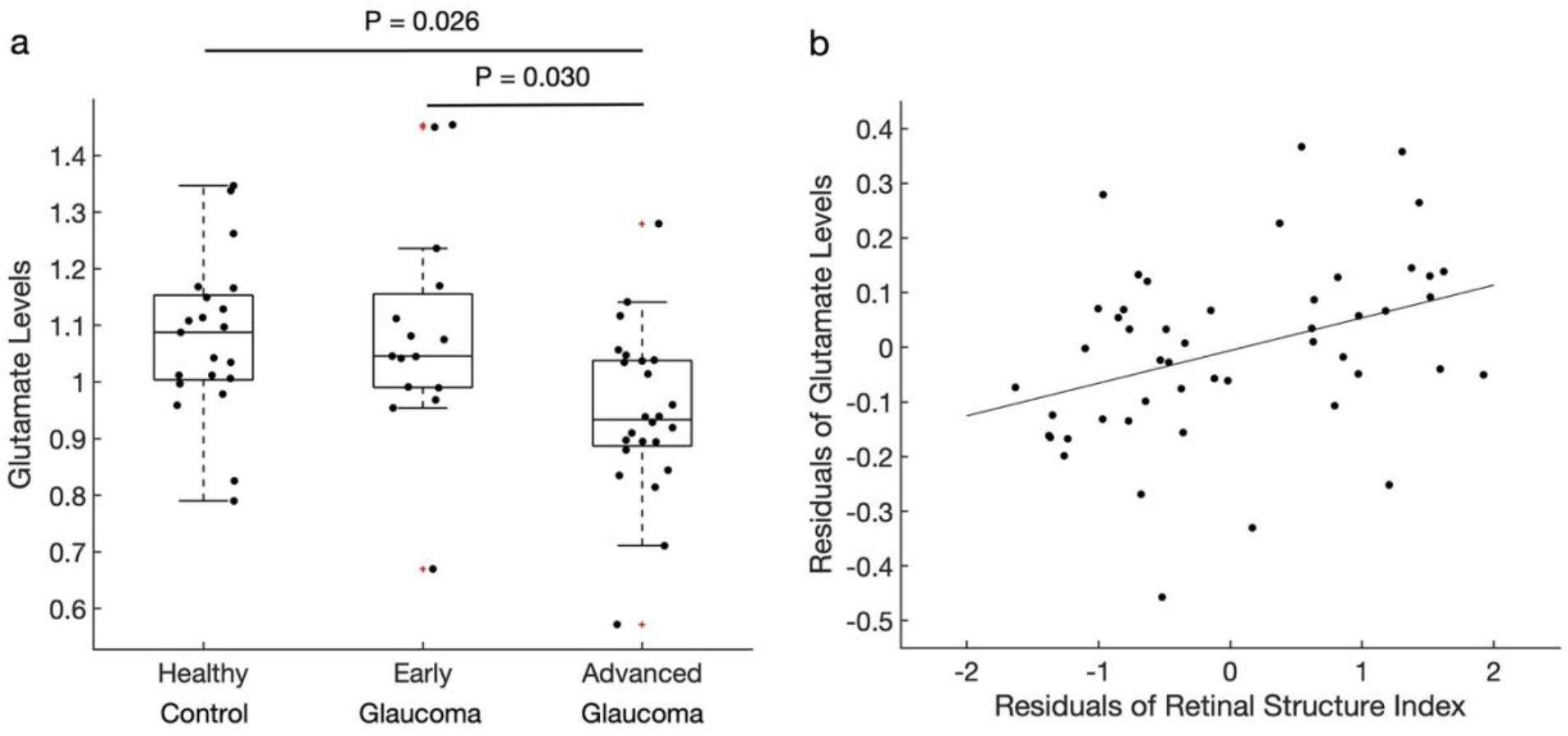
Glutamate levels in the visual cortex. **a,** The glutamate levels (normalized to total creatine levels) were significantly lower in advanced glaucoma patients compared to healthy controls (Bonferroni-corrected *P*=0.026) and early glaucoma patients (Bonferroni-corrected *P*=0.030). The distributions are represented using box plots and the outliers are plotted as red plus signs. Individual data points are overlaid on the box plots. **b,** The retinal structure index was positively correlated with the glutamate levels after controlling for age (r=0.364, *P*=0.011).

Given the significant reduction of GABA and glutamate in the visual cortex of glaucoma patients, we further tested whether the neural specificity is associated with the levels of GABA and glutamate (see **Extended Data Fig. 1** for box plots of the neural specificity). For this, we obtained each individual’s brain activity patterns across V1, V2, ventral V3, and ventral V4, which overlapped with the MRS voxel. During image acquisition, subjects viewed the horizontal and vertical flickering checkerboards in an alternating order. We measured the marker for neural specificity by calculating the fisher z-transformed correlation coefficient between the brain activation patterns representing horizontal and vertical flickering checkerboards. Here, more negative correlation coefficient value indicated a stronger neural specificity.

To investigate the relationship between neural specificity and the reduced levels of GABA and glutamate, we conducted a series of multiple linear regression analyses while controlling for the effect of age and retinal structure index. Here, we adjusted the effect of the retinal structure index in addition to age because the impairments in the retina may block some of the incoming visual signals, resulting in degraded neural specificity. Our regression analyses showed that those with lower GABA levels present weaker neural specificity in the visual cortex when the retinal structure index and age were adjusted (T(42)=-2.165, *P*=0.036, β=-0.317, partial correlation=-0.317, R^2^ change=0.088). This association between GABA and neural specificity survived even after we additionally controlled for the levels of glutamate (T(39)=-2.143, *P*=0.038, β=-0.328, partial correlation=-0.325, R^2^ change=0.091) and the gray matter volume of the corresponding visual areas (T(38)=-2.159, *P*=0.037, β=-0.334, partial correlation=-0.331, R^2^ change=0.094; **Fig. 4a**). In contrast, other factors such as the glutamate levels (T(38)=0.439, *P*=0.663, β=0.070, partial correlation=0.071, R^2^ change=0.004; **Fig. 4b**), age (T(38)=1.508, *P*=0.140, β=0.223, partial correlation=0.238, R^2^ change=0.046; **Extended Data Fig. 2a**), retinal structure index (T(38)=-1.690, *P*=0.099, β=-0.267, partial correlation=-0.264, R^2^ change=0.057; **Extended Data Fig. 2b**), and the gray matter volume of visual areas (T(38)=0.684, *P*=0.498, β=0.100, partial correlation=0.110, R^2^ change=0.009; **Extended Data Fig. 2c**) failed to account for the neural specificity. These results suggest that the most important predictor of neural specificity was the GABA levels in the visual cortex.

**Fig. 4:**
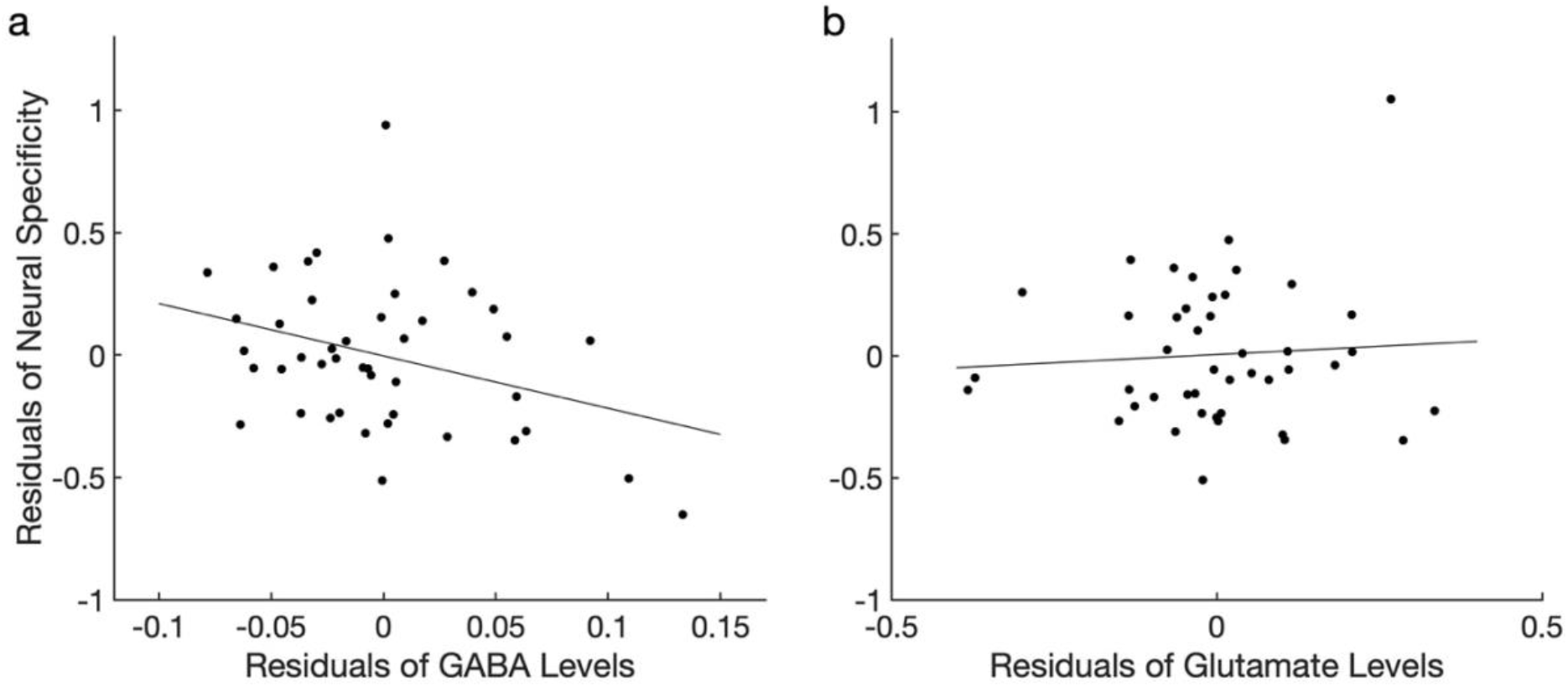
The relationship between neural specificity, GABA, and glutamate in the visual cortex for the entire sample. **a,** The GABA levels were significantly correlated with neural specificity after controlling for the glutamate levels, retinal structure index, age, and the gray matter volume of the visual areas (r=-0.331, *P*=0.037). **b,** The glutamate levels were not significantly associated with neural specificity after adjusting the effects of the GABA levels, retinal structure index, age, and the gray matter volume of the visual areas (r=0.110, *P*=0.498).

In a separate analysis, we conducted a hierarchical regression analysis to test whether the GABA levels explained the significant variance beyond that explained by the retinal structure index, age, glutamate levels, and the gray matter volume of visual areas. The results showed that adding GABA levels to the regression model explained an additional 9.4% of the variations in neural specificity and that this change was significant (F(1,38)=4.663, *P*=0.037).

After confirming that GABA levels significantly accounted for the neural specificity in the entire sample, we further tested whether the same association could be found within the samples of glaucoma patients and healthy controls separately. In the sample of glaucoma patients, we replicated the results of an association between GABA levels and neural specificity when the retinal structure index and age were controlled (T(27)=-2.561, *P*=0.016, β=-0.410, partial correlation=-0.442, R^2^ change=0.162). This association even became stronger when we additionally adjusted the effects of the levels of glutamate (T(25)=-2.929, *P*=0.007, β=-0.506, partial correlation=-0.506, R^2^ change=0.209) and the gray matter volume (T(24)=-2.907, *P*=0.008, β=-0.511, partial correlation=-0.510, R^2^ change=0.213; **Fig. 5a**). A hierarchical regression analysis also showed that an additional 21.3% of the variations in neural specificity can be explained by adding GABA levels to the regression model (F(1,24)=8.451, *P*=0.008).

**Fig. 5:**
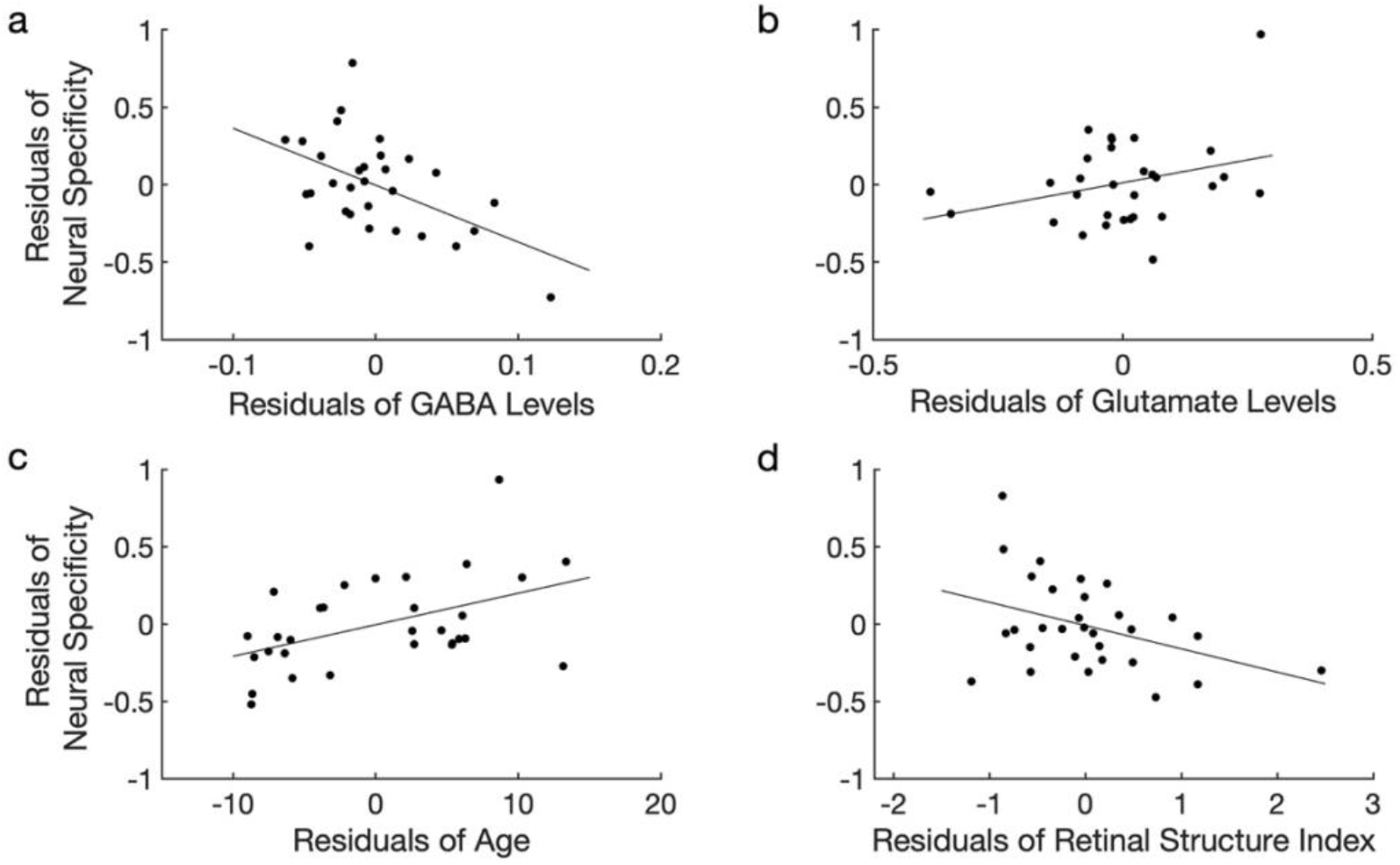
The relationship between neural specificity, GABA, glutamate, age, and the retinal structure index within the glaucoma group. **a,** The GABA levels significantly predicted neural specificity after controlling for the glutamate levels, retinal structure index, age, and the gray matter volume (r=-0.510, *P*=0.008). **b,** The glutamate levels were not significantly associated with neural specificity after controlling for the GABA levels, retinal structure index, age, and the gray matter volume (r=0.312, *P*=0.120). **c,** Age was significantly correlated with neural specificity after adjusting the effects of the GABA levels, glutamate levels, retinal structure index, and the gray matter volume (r=0.475, *P*=0.014). **d,** The retinal structure index was significantly associated with neural specificity after controlling for the GABA levels, glutamate levels, age, and the gray matter volume (r=-0.394, *P*=0.047).

Nevertheless, the glutamate levels failed to predict the neural specificity while controlling for the retinal structure index and age (T(29)=0.742, *P*=0.464, β=0.130, partial correlation=0.136, R^2^ change=0.015). Such null effect of the glutamate on the neural specificity remained the same when we additionally controlled the GABA levels (T(25)=1.578, *P*=0.127, β=0.282, partial correlation=0.301, R^2^ change=0.061), and the gray matter volume of the visual areas (T(24)=1.610, *P*=0.120, β=0.299, partial correlation=0.312, R^2^ change=0.065; **Fig. 5b**). This non-significant effect of the glutamate levels in the glaucoma group is consistent with a null effect of the glutamate observed in the entire sample combining both glaucoma and healthy subjects.

A discrepancy between the sample of glaucoma patients and the entire sample was seen in the effects of age and retinal structure index on neural specificity. Glaucoma patients with older ages presented weaker neural specificity while controlling for other confounding effects including retinal structure index, GABA levels, glutamate levels, and the gray matter volume of the visual areas (T(24)=2.644, *P*=0.014, β=0.467, partial correlation=0.475, R^2^ change=0.176; **Fig. 5c**). Similarly, those who had greater impairments on the retina structure exhibited reduced neural specificity when adjusting other confounding effects including age, GABA levels, glutamate levels, and the gray matter volume of the visual areas (T(24)=-2.097, *P*=0.047, β=-0.349, partial correlation=-0.394, R^2^ change=0.111; **Fig. 5d**). These results indicate that age and retinal structure index were meaningful predictors for neural specificity within the sample of glaucoma patients, whereas they were not in the entire sample combining both glaucoma and healthy subjects. The remaining factor, gray matter volume of visual areas did not have any association with the neural specificity within the glaucoma group (T(24)=0.441, *P*=0.663, β=0.073, partial correlation=0.090, R^2^ change=0.005; **Extended Data Fig. 3**).

Finally, we examined whether the same patterns of relationship could be seen within the healthy control group. The results showed that none of GABA, glutamate, age, retinal structure index, or gray matter volume of the visual areas was a significant predictor for the neural specificity in healthy controls after adjusting the remaining factors (GABA: T(8)=-0.678, *P*=0.517, β=-0.187, partial correlation=-0.233, R^2^ change=0.032; Glutamate: T(8)=0.058, *P*=0.956, β=0.018, partial correlation=0.020, R^2^ change<0.001; Age: T(8)=-0.436, *P*=0.674, β=-0.119, partial correlation=-0.152, R^2^ change=0.013; Retinal structure index: T(8)=0.117, *P*=0.910, β=0.033, partial correlation=0.041, R^2^ change=0.001; Gray matter volume: T(8)=2.243, *P*=0.055, β=0.643, partial correlation=0.621, R^2^ change=0.351; **Extended Data Fig. 4**).

Critically, the overall MRS results yielded by using total creatine for normalization were maintained even when we used N -acetyl-aspartate (NAA), another standard reference resonance for normalization. Using NAA for normalization, we still observed a trend of GABA reduction with increasing severity of glaucoma (one-way ANCOVA with a factor of group controlling for age, main effect of group: F(2,54)=2.822, *P*=0.068, partial η^2^=0.095). The GABA level normalized by NAA was also marginally associated with the neural specificity after adjusting the effects of the retinal structure index, age, the levels of glutamate normalized by NAA, and the gray matter volume in the entire sample (T(38)=-1.890, *P*=0.066, β=-0.286, partial correlation=-0.293, R^2^ change=0.073). In the sample of glaucoma patients, this association between the GABA level normalized by NAA and the neural specificity became even stronger (T(24)=-2.658, *P*=0.014, β=-0.481, partial correlation=-0.477, R^2^ change=0.180).

## Discussion

In the current study, we demonstrated that both GABA and glutamate levels in the visual cortex decline with increasing glaucoma severity. This association between impairments of the retina structure, GABA, and glutamate was independent of age among our older adults. Further, we showed that the reduction of GABA, but not that of glutamate, was associated with the neural specificity in the visual cortex after controlling for confounding effects including age and retinal structural damage. These results provide clear evidence that glaucomatous neurodegeneration involves the reduction of GABA and glutamate levels in the visual cortex and that the reduction of GABA, but not that of glutamate, plays a critical role in degrading neural specificity.

The reduction of GABA levels in the visual cortex of glaucoma patients is consistent with the literature showing that GABA metabolism is a unique pathway significantly associated with glaucoma^50^. Furthermore, GABA is susceptible to the accumulation of amyloid β and tau^24–27,53^. Prior studies demonstrated that glaucoma involves accumulation of tau in the vitreous fluid^17^, retina^16,18^, lateral geniculate nucleus^19^ as well as deposits of amyloid β in the lateral geniculate nucleus and the primary visual cortex^19^. Notably, exposure to amyloid β and tau is known to impair the GABAergic systems^24–27,53^. In animal models of Alzheimer’s disease, accumulation of amyloid β led to decreases in the number^26^ and density of GABAergic neurons before loss of pyramidal cells^27^. Further, amyloid β was shown to weaken synaptic inhibition via downregulating GABA_A_ receptors^24^. In another animal model expressing high deposits of tau, a significant reduction was observed in the GABA-synthetic enzyme and GABAergic interneurons, which co-localized with tau markers^25^. This line of studies raises a possibility that glaucoma may share common pathogenic mechanisms with Alzheimer’s disease and that accumulation of amyloid β and tau in the visual pathway may promote loss of GABA neurotransmission in glaucoma patients.

In addition to the possible impact of amyloid β and tau on GABA, several other factors may contribute to GABA reduction. First, impaired visual input may facilitate GABA reduction. Visual deprivation was shown to trigger GABA reduction in the primary visual cortex^54^. Given that the visual signals are gradually blocked as glaucoma progresses, it is speculated that its impact on GABA would increase as well. Consistent with this, our study observed that the GABA levels in the visual cortex measured by MRS decrease with increasing impairments of the retina structure. Second, accelerated aging process in glaucoma may affect GABA levels. Facilitated aging in glaucoma is supported by the finding that glaucoma patients present elevated amount of advanced glycation end products, which are inevitable compounds of the aging process, as well as upregulation of their receptors in the retina and optic nerve head^43^. Relatedly, the aging process involves impairments of GABAergic systems^55^. In healthy animal models and humans, age-related declines were observed in terms of the number of GABAergic neurons^56–58^, baseline GABA levels, GABA release, GABA receptor binding^57^, and MRS GABA measures throughout the brain^44–49^. Therefore, it is possible that the age-related declines of GABA may be accelerated in glaucoma patients. Future studies could address this possibility by obtaining the measures of biological speed of aging from biological assays along with MRS and functional MRI.

In this study, MRS measures of GABA cannot distinguish whether the decreased GABA levels in the visual cortex of glaucoma patients reflect reduction of GABAergic neurons, GABA neurotransmitters, or GABA synthesis^59^. Any combination of these changes could lead to reduced MRS GABA levels. While it is yet unknown which factor contributed the most to our MRS observations, one study using an experimental monkey model observed that after unilateral IOP elevation, the GABA_A_ receptor protein in the primary visual cortex was reduced^11^, possibly driven by suppression of visually driven activity^60^. This finding suggests that the declined MRS GABA levels that we observed in the current study are in part reflective of reduced GABA receptor binding. Other possibilities such as reduction of the GABAergic neurons or GABA neurotransmitters need to be further identified in future *in vivo, in situ*, and *ex vivo* studies using a finer spatial resolution.

The decreased GABA signals in the visual cortex of glaucoma patients suggest that the cortical GABAergic function is likely impaired in glaucoma. GABAergic signals are critical for shaping the response of a single neuron as well as the response patterns of a population of neurons^28^. At the level of a single neuron, GABA-mediated inhibition can sharpen the neuronal response around the preferred stimulus feature^61^. In line with this, application of GABA agonist increases the neuron’s selectivity to preferred stimulus feature, while application of GABA antagonist decreases the neuron’s selectivity^62^. Therefore, it is speculated that the selectivity of neurons in the visual cortex may be undermined in glaucoma patients. At the population level, GABAergic signals are related to the specificity of neuronal activity patterns in response to different stimuli. With lower amount of GABA, neuronal activity patterns become less distinct or more confusable^29,30^. Consistent with these prior findings, we observed that those who had lower GABA levels displayed less specific neural activity patterns in the visual cortex independent of the age and impairments of the retina structure. It suggests that the declines of GABA and neural specificity may negatively affect the downstream processes along the lower-to higher-order brain regions^32,33^ in glaucoma.

Alterations of GABA levels and neural specificity in glaucoma patients are likely to have functional influences to various aspects of brain functions. Prior studies showed that GABA levels in healthy subjects are associated with visual-spatial intelligence, visual surround suppression^63^, motor inhibition^45^, motor learning^64^, general sensorimotor functioning^49^, cognitive performance^46,48^, experience-dependent plasticity^65^, and sleep^66^. Further, neural specificity was found to be related to memory performance^32^ and fluid processing ability^33^. This line of evidence suggests that these aspects of brain functions may be undermined if the GABA level and the neural specificity are impaired. In line with this, some studies reported abnormalities in perceptual, cognitive functions and sleep quality in glaucoma patients. For example, glaucoma patients were found to have elevated visual crowding effect^34^, impairments on perceptual switch during binocular rivalry^35–37^, declines in visual categorization^39,40^, visual attention^41^ and cognitive capacity^39,40,42^ as well as sleep disturbance^67,68^. Thus, we propose that there could be a functional link between GABA reduction and the reported abnormalities in glaucoma patients. Future studies could address this question by employing both MRS and behavioral measures.

In addition to the GABA levels, we observed that the impairments on the retina structure and the years of age could predict the neural specificity within the sample of glaucoma patients. Specifically, lower neural specificity was associated with greater impairments on the retina structure and older ages. These associations are reasonable given that impaired retina structure blocks parts of the incoming visual signals and aging is accompanied by degraded neural distinctiveness/discriminability^29–31,69,70^. However, these relationships between GABA, retinal structure index, years of age and neural specificity were not observed within our sample of healthy subjects. This discrepancy could be partly due to the fact that healthy subjects had relatively less affected GABA, retinal structure, and neural specificity compared to glaucoma patients, making the association between these factors less obvious. Further, the small sample size and the limited age range could have made it difficult to detect any meaningful association in healthy subjects. However, it should also be noted that such discrepancy between glaucoma and healthy subjects, which is that the neural specificity can be explained by the retinal structure and the years of age in glaucoma patients, but not in the healthy controls, may be in part associated with accelerated aging process in glaucoma patients.

Another important finding in this study is that the glutamate levels in the visual cortex decline with the impairments on the retinal structure. This observation may appear inconsistent with the literature showing that accumulation of amyloid β leads to increased glutamate levels in the extracellular space^20,21^. Nevertheless, it should be noted that glutamate and GABA are inseparable signals as GABA is synthesized by decarboxylation of glutamate via glutamate acid decarboxylase. Increases or decreases in the strength of cortical excitation are known to be accompanied by the corresponding proportional changes in the strength of inhibition^71,72^. Further, when cortical excitation or inhibition is acutely manipulated, the neurons become dysfunctional^73^. Thus, our observation of reduced glutamate levels in the visual cortex may be at least partially related to the reduced GABA levels. Further, recent studies have demonstrated that amyloid β and glutamatergic signals have multifaceted mechanisms^74^. In the early stage of Alzheimer’s disease, glutamatergic activity is elevated leading to hyperexcitability of neurons^75,76^. However, as the Alzheimer’s disease progresses, glutamate release and vesicular glutamate transporters become markedly decreased^77^. Prior MRS studies also observed a decreasing pattern of glutamate levels with severity of Alzheimer’s disease^78,79^. Thus, our results of decreased glutamate levels in glaucoma may resemble these progressive glutamate changes in the late phases of Alzheimer’s disease.

While the current study focused on the visual cortex only, it is worth knowing whether the same neurochemical changes occur in the retina of glaucoma patients. A number of patient and experimental animal studies have examined the glutamate-mediated excitotoxicity in the vitreous body and retina in glaucoma^80–83^. Nevertheless, these studies provided disparate results, making it unclear whether the glutamate in the retina is elevated under glaucoma. Other studies investigating GABA, however, point to a dysfunction of the GABAergic signals in glaucoma. For example, in animal models of glaucoma, the retina showed decreases in the GABA turnover rate, activity of GABA-synthetic enzyme, GABA release and the expression of retinal genes associated with GABAergic systems^51,84^. Further, studies reported that modulating GABAergic activity in the rat’s retina under chronic glaucomatous conditions may improve retinal ganglion cell viability and function^85,86^. Future studies are warranted to examine whether GABA plays a role in glaucomatous neuroprotection and functional recovery throughout the visual system of the brain.

To conclude, our findings demonstrate a tight link between declines of GABA and neural specificity in the visual cortex of glaucoma patients. Our results suggest that glaucoma-specific declines of GABA undermine neural specificity in the visual cortex and that targeting GABA could improve the neural specificity in glaucoma patients.

## Methods

### Subjects

The Institutional Review Board of New York University Grossman School of Medicine approved this study. This study adhered to the tenets of the Declaration of Helsinki. Informed consent was obtained from all subjects prior to participation. Forty-one glaucoma subjects [age= 65.88 ± 1.23 (mean ± S.E.M.); 41.5% male] and twenty-three healthy controls [age= 64.78 ± 1.63 (mean ± S.E.M.); 47.8% male] were included in the study between August 2018 and May 2022 at New York University Langone Health’s Department of Ophthalmology. All subjects had a best corrected visual acuity (BCVA) of 20/60 or better and showed no past medical history (PMH) or current evidence of retinal or neurological disorders. The demographic and clinical information of those who were included in this study is depicted in **Table 1**. Subjects in the disease group were clinically diagnosed with primary glaucoma, whereas healthy controls exhibited no clinical evidence of glaucomatous conditions. Subjects were not allowed to participate in the MRI study if they were pregnant or breastfeeding at the time of the study, had any metal parts or fragments in the body with the exception of dental fillings, had conditions such as anxiety or claustrophobia, or had obesity that would hinder placement into the MRI scanner.

### Clinical ophthalmic exams

The clinical ophthalmic data for both glaucoma patients and healthy subjects were collected, which included pRNFL thickness, mGCIPL thickness, optic nerve head C/D ratio, and NRR area through an automatic analytic software on board of the Cirrus spectral-domain optical coherence tomography device (Zeiss, Dublin, CA, USA). Visual field MD was also obtained from the Humphrey Swedish Interactive Thresholding Algorithm (SITA) 24-2 standard (Zeiss, Dublin, CA, USA). The average MD of the left and right eyes combined (OU) were obtained to assign the patients into early or advanced glaucoma group. Early glaucoma was categorized as glaucoma patients with average OU MD better than −6.0dB and advanced glaucoma was determined as average OU MD worse than −6.0dB.

### MRI data acquisition

Age-matched healthy subjects and glaucoma patients were scanned inside a 3-Tesla MR Prisma scanner (Siemens, Germany) with a 20-channel head coil at the Center for Biomedical Imaging, NYU Langone health, New York University. For anatomical reconstruction, high-resolution T1-weighted MR images were acquired using a multi-echo magnetization-prepared rapid gradient echo sequence, with 256 slices, voxel size=0.8 × 0.8 × 0.8 mm, 0-mm slice gap, repetition time (TR)=2400 ms, echo time (TE)=2.24 ms, flip angle=8°, field of view= 256 mm, and bandwidth=210 Hz per pixel.

For MRS acquisition, we manually positioned a 2.2 × 2.2 × 2.2 cm^3^ voxel along the calcarine sulci in the occipital cortex. The voxel covered parts of the visual cortex including V1, V2, V3, and V4. Shimming was performed by a vendor-provided automated shim tool followed by manual fine adjustment. The shim value defined by the full width at half maximum of the water peak was 13.02 ± 0.13 (mean ± S.E.M.) for gamma-aminobutyric acid (GABA) and 12.93 ± 0.14 (mean ± S.E.M.) for glutamate. The concentration of GABA was measured from a voxel using a MEshcher-GArwood-Point RESolved Spectroscopy (MEGA-PRESS) sequence with double-banded pulses, at TR=1500 ms, TE=68 ms, number of average=172, and scanning duration=522 s. The final spectrum was obtained by subtracting the ‘edit-off’ spectrum from ‘edit-on’ spectrum. The concentration of glutamate was measured from the same voxel using a Point RESolved Spectroscopy (PRESS) sequence, with TR=3000 ms, TE=30 ms, number of average=99, and scanning duration=300 s. During both MEGA-PRESS and PRESS scans, subjects performed a fixation task. For two subjects, MRS acquisition could not be completed due to technical issues.

After MRS acquisition, functional MR images were acquired using a gradient-echo echo-planar imaging (EPI) sequence, with voxel size=2.3 × 2.3 × 2.3 mm, TR=1000 ms, TE=32.60 ms, and scanning duration=300 s. During fMRI scan, subjects performed a fixation task while a flickering checkerboard pattern was presented at the horizontal vs. vertical meridians. For four subjects, fMRI images could not be obtained due to technical issues.

### Fixation task

To maintain subjects’ attention and vigilance levels across the scans, we provided subjects with the fixation task during MEGA-PRESS and PRESS scans. The fixation point was presented at the center of the gray background and subjects were asked to fixate their eyes at the point. The fixation point changed its color unpredictably from white ([R, G, B]=[255, 255, 255] to red ([R, G, B]=[255, 127, 127]) and returned to white 1.5 s later. When the button was pressed within a 1.5-s time window, this response was recorded as a hit. If not, the response was recorded as a miss. The mean accuracy (± S.E.M.) was 98.87 ± 0.58%. The accuracy did not differ across healthy controls, and early and advanced glaucoma patients (F(2,32)=0.337, *P*=0.716, partial η^2^=0.021).

### Neural specificity task

During fMRI scans (1 run=300 s), subjects performed the same fixation task, while the background was filled with flickering checkerboard patterns. The checkerboard patterns were presented at either horizontal or vertical meridians in a gray background. The purpose of adding the flickering checkerboard patterns in the background was to obtain the neural specificity for horizontal and vertical meridians. Each of the horizontal and vertical meridians was presented for 8 s in alternation, with 18 trials for horizontal meridians and 18 trials for vertical meridians in total. At the first and the last 6-s periods, the fixation point was presented only without any checkerboard patterns. The mean accuracy of the fixation task (± S.E.M.) was 94.10 ± 1.11% and was comparable across healthy controls, and early and advanced glaucoma patients (F(2,42)=1.529, *P*=0.229, partial η^2^=0.068).

### MRS data analysis

MRS data was fitted in the frequency domain using the LCModel software^87^. We assessed the quality of fitting by visually inspecting LCmodel fitted spectra and examining their Cramer-Rao lower bounds (CRLB) and spectral signal-to-noise (S/N) ratio. Four spectra from the MEGA-PRESS and two spectra from the PRESS scans were excluded from further analyses because of poor fitting (CRLB>20%, S/N<8). The CRLB was 8.76 ± 0.15% (mean ± S.E.M.) for GABA and 8.65 ± 0.40% (mean ± S.E.M.) for glutamate. The S/N ratio was 23.84 ± 0.38 (mean ± s.e.m.) for GABA and 26.6 ± 1.36 (mean ± s.e.m.) for glutamate. We normalized the concentrations of GABA and glutamate using the amount of total creatine, which is commonly used as a standard reference resonance^87,88^. For complementary support, we also normalized the GABA and glutamate concentrations by the amount of N-acetyl-aspartate (NAA).

### fMRI data analysis

We analyzed fMRI data using Freesurfer software (http://surfer.nmr.mgh.harvard.edu/) and MATLAB. The fMRI data was preprocessed with motion correction but not with spatial or temporal smoothing. Then the functional data was registered to the individual’s structural template. We extracted the blood-oxygenation-level-dependent (BOLD) signals from the individual’s region-of-interest (ROI) masks generated from Freesurfer’s cortical reconstruction process. The ROI masks that we used were four cytoarchitectonic areas in the occipital lobe (hOc1-hOc4v) which spatially overlapped with the location of the MRS voxel. The extracted BOLD signals were then shifted by 6 s to account for the hemodynamic delay. We excluded voxels with spikes greater than 10 standard deviations from the mean and removed the linear trend in the BOLD time course. Then we normalized the BOLD signals using z-score for each voxel.

### Calculation of neural specificity

We defined the neural specificity as the dissimilarity in brain responses between different visual conditions. For this, we averaged the z-normalized BOLD signals of each voxel across 8 volumes (8 s) which corresponded to the duration of horizontal/vertical meridian presentation. These BOLD signals of each voxel were then averaged across 18 trials of horizontal meridians or 18 trials of vertical meridians separately. Given that these two patterns of BOLD signals represented the horizontal or vertical meridians, we computed the Fisher z-transformed correlation coefficient between these two representations. Finally, we averaged the correlation coefficient for each representation across V1, V2, ventral V3, and ventral V4.

### Statistics

The sample size was not predetermined but similar or even greater to those reported in prior studies^2,3,10,12,13,89^. The investigators were not blinded to disease diagnosis during data collection. For all statistical comparisons, we conducted two-tailed parametric tests with P<0.05 as the criterion for statistical significance. For ANOVAs, we assessed the assumption of homogeneity of variances using Levene’s test. The following post-hoc tests were conducted with Bonferroni corrections. For multiple regression analyses, we confirmed that the assumptions of independence of observations and non-multicollinearity were not violated using the Durbin-Watson statistic and tolerance/VIF values. Further, we extracted a common component from clinical ophthalmic measures including pRNFL thickness, mGCIPL thickness, optic nerve head C/D ratio, and NRR area for each individual using PCA. Then we entered this component, a measure of retina structure in the regression model rather than entering all clinical ophthalmic measures separately because the clinical measures were highly correlated to each other, violating the non-multicollinearity. We set the criteria of selecting the component in the PCA as the eigenvalue higher than one.

### Apparatus

We created all visual stimuli in MATLAB using Psychophysics Toolbox 3^90^. The stimuli were presented via an MRI-compatible projector (1024 × 768 resolution, 60 Hz refresh rate).

## Data and code availability

Data and codes for analysis are freely available at https://osf.io/85cds/

## Acknowledgements

We would like to thank Ms Tonya Robins, Zena Moore, Jamika Singleton-Garvin and members of the Neuroimaging and Visual Science Laboratory at New York University Grossman School of Medicine for their help with subject recruitment and technical support. This work is supported in part by the National Institutes of Health R01-EY028125, R01-EY013178, and P41-EB017183 (Bethesda, Maryland), BrightFocus Foundation G2016030, G2019103, and G2021001F (Clarksburg, Maryland), and an unrestricted grant from Research to Prevent Blindness to NYU Langone Health Department of Ophthalmology (New York, New York).

---

**Extended Data Fig. 1:**
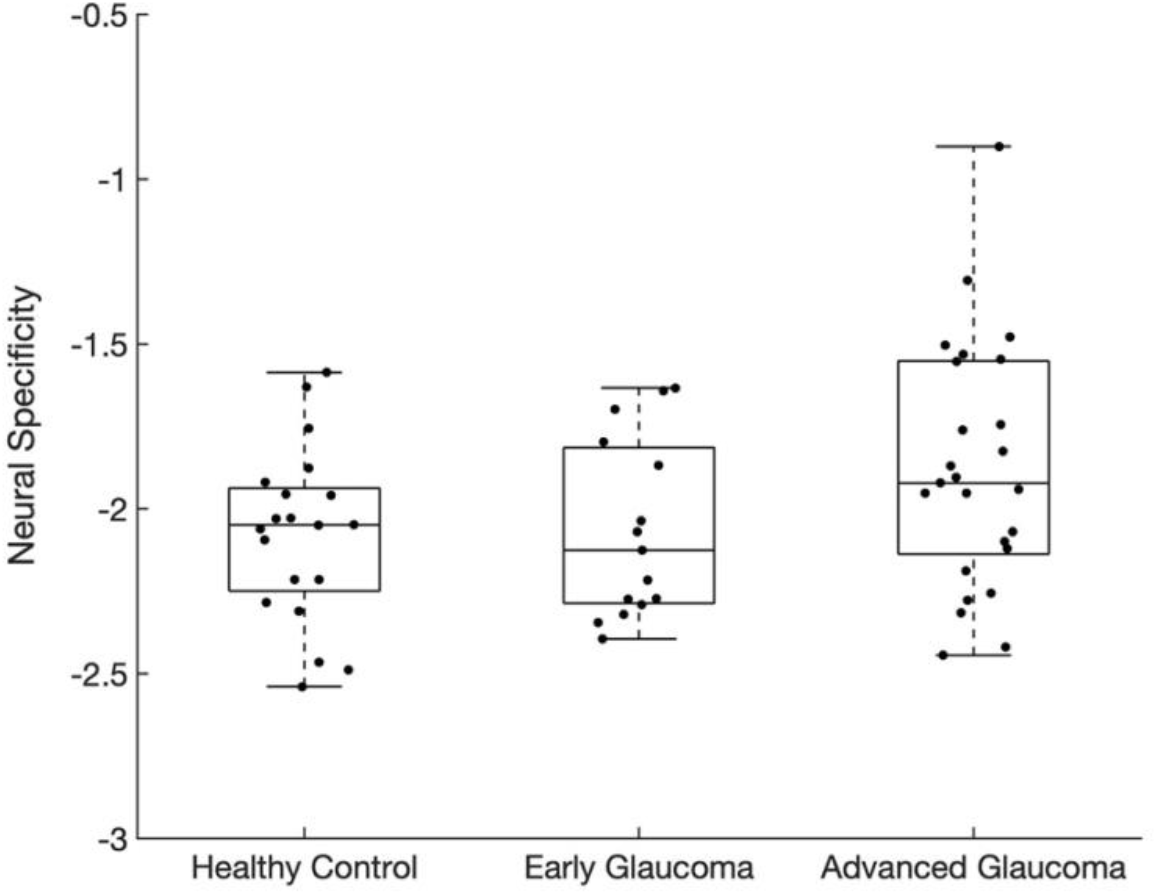
Neural specificity across groups. The group effect was not significant, although there was a trend toward significance (F(2,56)=2.595, *P*=0.084, partial η^2^=0.085). The distributions are represented using box and whisker plots.

**Extended Data Fig. 2:**
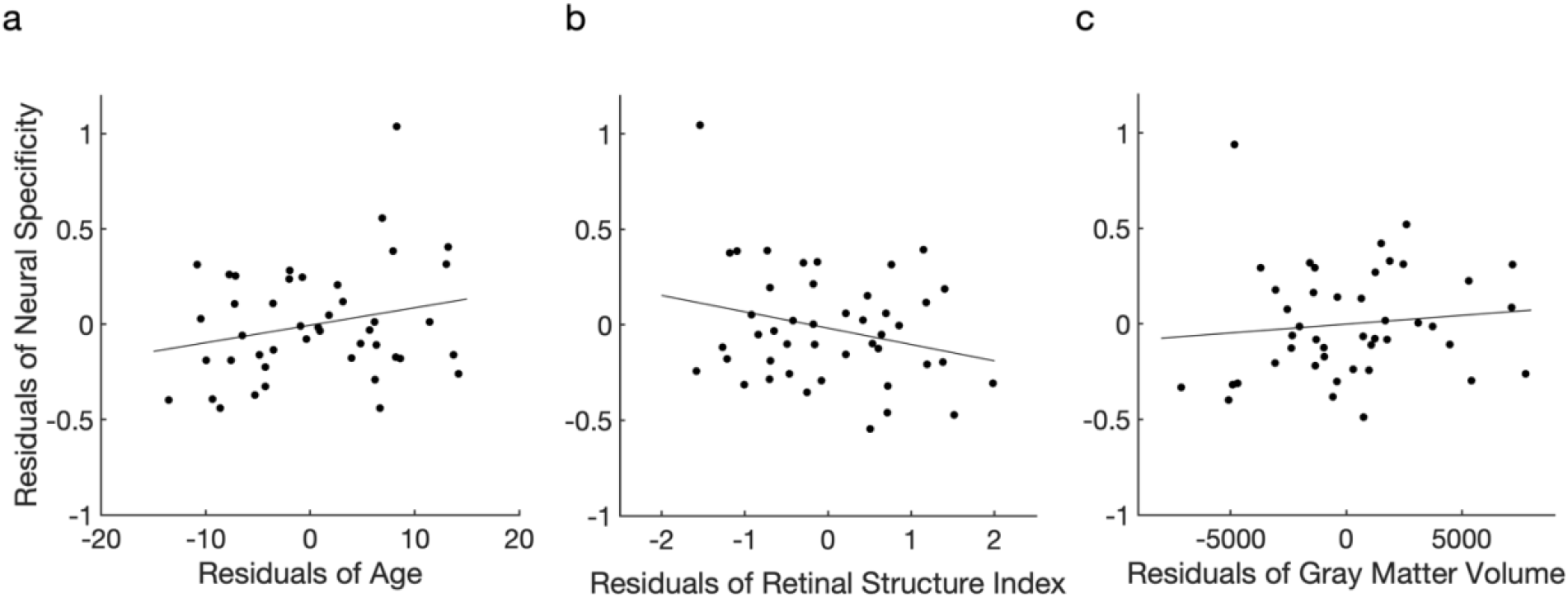
The relationship between neural specificity, age, retinal structure, and gray matter volume in the entire sample including glaucoma and healthy subjects. **a,** Age was not significantly associated with neural specificity after adjusting the effects of the GABA, glutamate, retinal structure index, and the gray matter volume (r=0.238, *P*=0.140). **b,** The retinal structure index did not significantly predict the neural specificity after controlling for the GABA, glutamate, age, and the gray matter volume (r=-0.264, *P*=0.099). **c,** The gray matter volume of the visual cortex was not significantly correlated with the nerual specificity after adjusting the effects of the GABA, glutamate, retinal structure index, and age (r=0.110, *P*=0.498).

**Extended Data Fig. 3:**
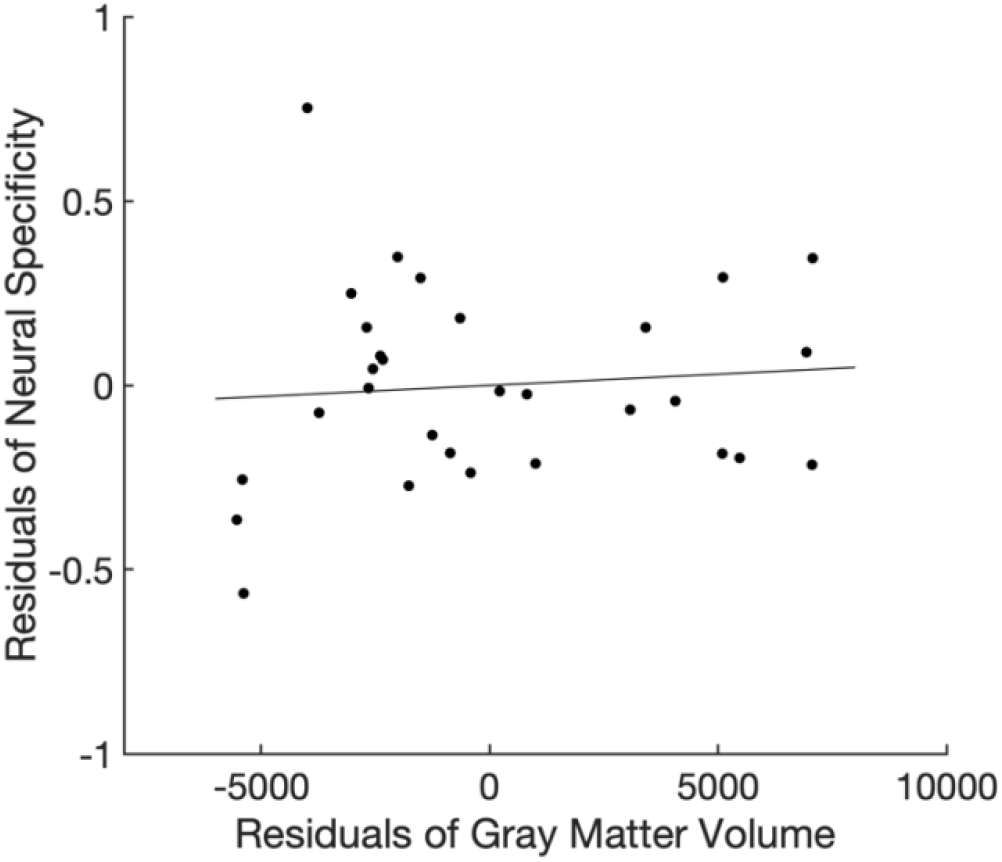
The relationship between neural specificity and gray matter volume within the glaucoma group. The gray matter volume was not correlated with neural specificity (r=0.090, *P*=0.663).

**Extended Data Fig. 4:**
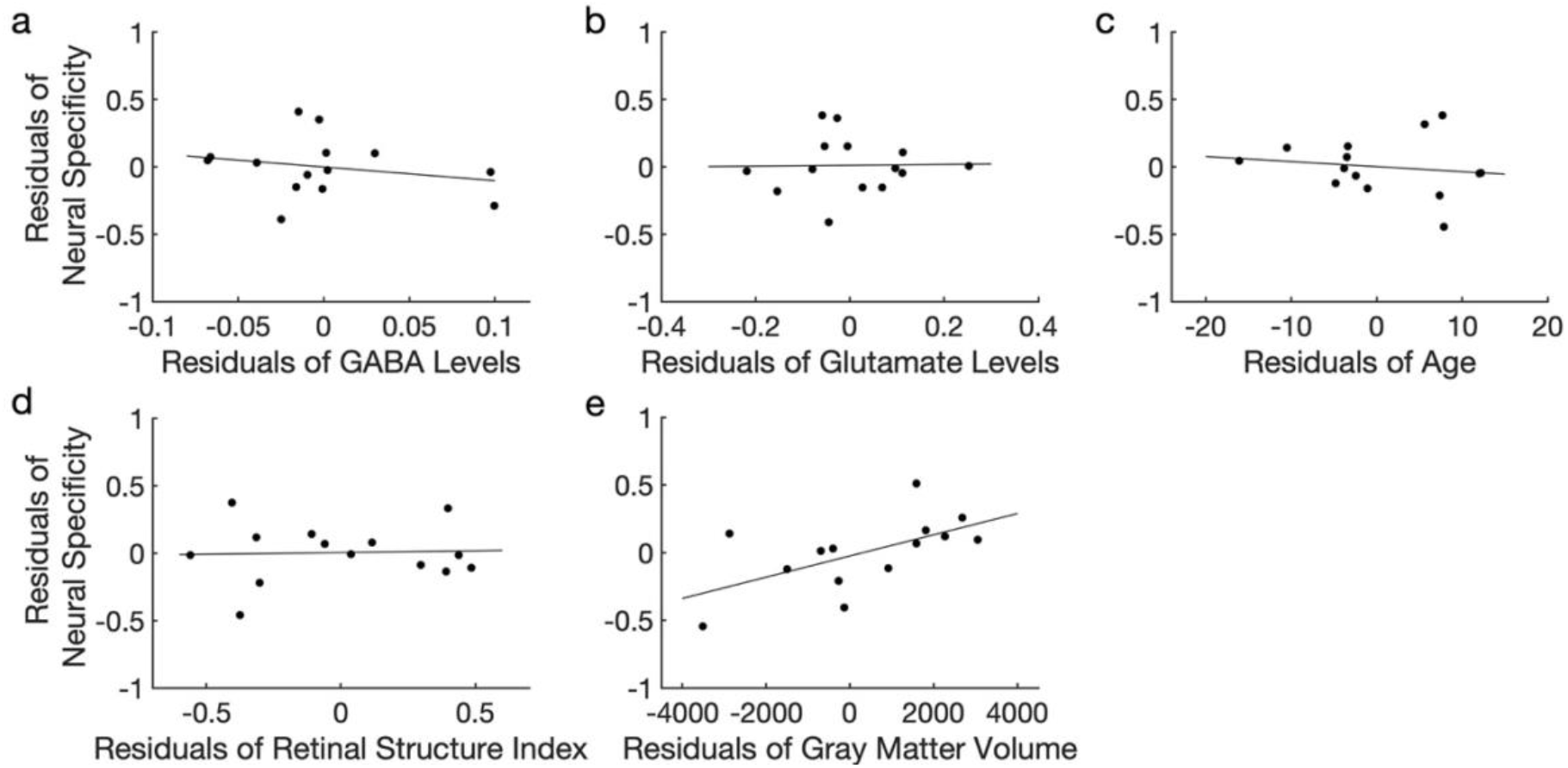
The relationship between neural specificity, GABA, and glutamate in the sample of healthy subjects. **a,** The GABA levels were not associated with neural specificity after adjusting the effects of the glutamate, retina structure, age, and the gray matter volume (r=-0.233, *P*=0.517). **b,** The glutamate did not significantly predict neural specificity after controlling the effects of the GABA, retina structure, age, and the gray matter volume (r=0.020, *P*=0.956). **c,** The age was not correlated with neural specificity after adjusting the effects of the GABA, glutamate, retina structure, and the gray matter volume (r=-0.152, *P*=0.674). **d,** The retina structure was not associated with neural specificity after controlling for the effects of the GABA, glutamate, age, and the gray matter volume (r=0.041, *P*=0.910). **e,** The gray matter volume did not significantly predict neural specificity after adjusting the effects of the GABA, glutamate, retina structure, and age (r=0.621, *P*=0.055).

## Notes

https://osf.io/85cds/

### Competing Interest Statement

J.S.S. has made the following disclosure: Royalties e Zeiss, Dublin, CA (for intellectual property licensed by the Massachusetts Institute of Technology and Massachusetts Eye and Ear Infirmary).

### Summary of Updates

Abstract revised.

## References

1. Tham, Y.C., et al. Global prevalence of glaucoma and projections of glaucoma burden through 2040: a systematic review and meta-analysis. Ophthalmology 121, 2081–2090 (2014).

2. Garaci, F.G., et al. Optic nerve and optic radiation neurodegeneration in patients with glaucoma: in vivo analysis with 3-T diffusion-tensor MR imaging. Radiology 252, 496–501 (2009).

3. Zikou, A.K., et al. Voxel-based morphometry and diffusion tensor imaging of the optic pathway in primary open-angle glaucoma: a preliminary study. AJNR Am J Neuroradiol 33, 128–134 (2012).

4. Weber, A.J., Chen, H., Hubbard, W.C. & Kaufman, P.L. Experimental glaucoma and cell size, density, and number in the primate lateral geniculate nucleus. Invest Ophthalmol Vis Sci 41, 1370–1379 (2000).

5. Yucel, Y.H., Zhang, Q., Gupta, N., Kaufman, P.L. & Weinreb, R.N. Loss of neurons in magnocellular and parvocellular layers of the lateral geniculate nucleus in glaucoma. Arch Ophthalmol 118, 378–384 (2000).

6. Crawford, M.L., Harwerth, R.S., Smith, E.L., 3rd, Shen, F. & Carter-Dawson, L. Glaucoma in primates: cytochrome oxidase reactivity in parvo- and magnocellular pathways. Invest Ophthalmol Vis Sci 41, 1791–1802 (2000).

7. Imamura, K., et al. Molecular imaging reveals unique degenerative changes in experimental glaucoma. Neuroreport 20, 139–144 (2009).

8. Chaturvedi, N., Hedley-Whyte, E.T. & Dreyer, E.B. Lateral geniculate nucleus in glaucoma. Am J Ophthalmol 116, 182–188 (1993).

9. Gupta, N., Ang, L.C., Noel de Tilly, L., Bidaisee, L. & Yucel, Y.H. Human glaucoma and neural degeneration in intracranial optic nerve, lateral geniculate nucleus, and visual cortex. Br J Ophthalmol 90, 674–678 (2006).

10. Engelhorn, T., et al. Diffusion tensor imaging detects rarefaction of optic radiation in glaucoma patients. Acad Radiol 18, 764–769 (2011).

11. Lam, D.Y., Kaufman, P.L., Gabelt, B.T., To, E.C. & Matsubara, J.A. Neurochemical correlates of cortical plasticity after unilateral elevated intraocular pressure in a primate model of glaucoma. Invest Ophthalmol Vis Sci 44, 2573–2581 (2003).

12. Boucard, C.C., et al. Changes in cortical grey matter density associated with long-standing retinal visual field defects. Brain 132, 1898–1906 (2009).

13. Murphy, M.C., et al. Retinal Structures and Visual Cortex Activity are Impaired Prior to Clinical Vision Loss in Glaucoma. Sci Rep-Uk 6(2016).

14. Ghiso, J.A., Doudevski, I., Ritch, R. & Rostagno, A.A. Alzheimer’s disease and glaucoma: mechanistic similarities and differences. J Glaucoma 22 Suppl 5, S36–38 (2013).

15. Sen, S., Saxena, R., Tripathi, M., Vibha, D. & Dhiman, R. Neurodegeneration in Alzheimer’s disease and glaucoma: overlaps and missing links. Eye (Lond) 34, 1546–1553 (2020).

16. Gupta, N., Fong, J., Ang, L.C. & Yucel, Y.H. Retinal tau pathology in human glaucomas. Can J Ophthalmol 43, 53–60 (2008).

17. Yoneda, S., et al. Vitreous fluid levels of beta-amyloid((1-42)) and tau in patients with retinal diseases. Jpn J Ophthalmol 49, 106–108 (2005).

18. Chiasseu, M., et al. Tau Accumulation, Altered Phosphorylation, and Missorting Promote Neurodegeneration in Glaucoma. J Neurosci 36, 5785–5798 (2016).

19. Yan, Z., et al. Elevated Intraocular Pressure Induces Amyloid-beta Deposition and Tauopathy in the Lateral Geniculate Nucleus in a Monkey Model of Glaucoma. Invest Ophthalmol Vis Sci 58, 5434–5443 (2017).

20. Fernandez-Tome, P., Brera, B., Arevalo, M.A. & de Ceballos, M.L. Beta-amyloid25-35 inhibits glutamate uptake in cultured neurons and astrocytes: modulation of uptake as a survival mechanism. Neurobiol Dis 15, 580–589 (2004).

21. Harris, M.E., et al. Amyloid beta peptide (25-35) inhibits Na+-dependent glutamate uptake in rat hippocampal astrocyte cultures. J Neurochem 67, 277–286 (1996).

22. Chin, J.H., Ma, L., MacTavish, D. & Jhamandas, J.H. Amyloid beta protein modulates glutamate-mediated neurotransmission in the rat basal forebrain: involvement of presynaptic neuronal nicotinic acetylcholine and metabotropic glutamate receptors. J Neurosci 27, 9262–9269 (2007).

23. Kabogo, D., Rauw, G., Amritraj, A., Baker, G. & Kar, S. ss-amyloid-related peptides potentiate K+-evoked glutamate release from adult rat hippocampal slices. Neurobiol Aging 31, 1164–1172 (2010).

24. Ulrich, D. Amyloid-beta Impairs Synaptic Inhibition via GABA(A) Receptor Endocytosis. J Neurosci 35, 9205–9210 (2015).

25. Levenga, J., et al. Tau pathology induces loss of GABAergic interneurons leading to altered synaptic plasticity and behavioral impairments. Acta Neuropathol Commun 1, 34 (2013).

26. Krantic, S., et al. Hippocampal GABAergic neurons are susceptible to amyloid-beta toxicity in vitro and are decreased in number in the Alzheimer’s disease TgCRND8 mouse model. J Alzheimers Dis 29, 293–308 (2012).

27. Ramos, B., et al. Early neuropathology of somatostatin/NPY GABAergic cells in the hippocampus of a PS1xAPP transgenic model of Alzheimer’s disease. Neurobiol Aging 27, 1658–1672 (2006).

28. Isaacson, J.S. & Scanziani, M. How inhibition shapes cortical activity. Neuron 72, 231–243 (2011).

29. Chamberlain, J.D., et al. GABA levels in ventral visual cortex decline with age and are associated with neural distinctiveness. Neurobiol Aging 102, 170–177 (2021).

30. Lalwani, P., et al. Neural distinctiveness declines with age in auditory cortex and is associated with auditory GABA levels. Neuroimage 201, 116033 (2019).

31. Simmonite, M. & Polk, T.A. Age-related declines in neural distinctiveness correlate across brain areas and result from both decreased reliability and increased confusability. Neuropsychol Dev Cogn B Aging Neuropsychol Cogn 29, 483–499 (2022).

32. Fandakova, Y., Leckey, S., Driver, C.C., Bunge, S.A. & Ghetti, S. Neural specificity of scene representations is related to memory performance in childhood. Neuroimage 199, 105–113 (2019).

33. Park, J., Carp, J., Hebrank, A., Park, D.C. & Polk, T.A. Neural specificity predicts fluid processing ability in older adults. J Neurosci 30, 9253–9259 (2010).

34. Ogata, N.G., et al. Visual Crowding in Glaucoma. Invest Ophthalmol Vis Sci 60, 538–543 (2019).

35. Tarita-Nistor, L., Samet, S., Trope, G.E. & Gonzalez, E.G. Intra- and inter-hemispheric processing during binocular rivalry in mild glaucoma. PLoS One 15, e0229168 (2020).

36. Issashar Leibovitzh, G., Trope, G.E., Buys, Y.M. & Tarita-Nistor, L. Perceptual Grouping During Binocular Rivalry in Mild Glaucoma. Front Aging Neurosci 14, 833150 (2022).

37. Joao, C.A.R., Scanferla, L. & Jansonius, N.M. Binocular Interactions in Glaucoma Patients With Nonoverlapping Visual Field Defects: Contrast Summation, Rivalry, and Phase Combination. Invest Ophthalmol Vis Sci 62, 9 (2021).

38. Tarita-Nistor, L., Samet, S., Trope, G.E. & Gonzalez, E.G. Dominance wave propagation during binocular rivalry in mild glaucoma. Vision Res 165, 64–71 (2019).

39. Lenoble, Q., Lek, J.J. & McKendrick, A.M. Visual object categorisation in people with glaucoma. Br J Ophthalmol 100, 1585–1590 (2016).

40. Roux-Sibilon, A., et al. Scene and human face recognition in the central vision of patients with glaucoma. PLoS One 13, e0193465 (2018).

41. Swenor, B.K., et al. Impact of the Ability to Divide Attention on Reading Performance in Glaucoma. Invest Ophthalmol Vis Sci 58, 2456–2462 (2017).

42. Gangeddula, V., Ranchet, M., Akinwuntan, A.E., Bollinger, K. & Devos, H. Effect of Cognitive Demand on Functional Visual Field Performance in Senior Drivers with Glaucoma. Front Aging Neurosci 9, 286 (2017).

43. Tezel, G., Luo, C. & Yang, X. Accelerated aging in glaucoma: immunohistochemical assessment of advanced glycation end products in the human retina and optic nerve head. Invest Ophthalmol Vis Sci 48, 1201–1211 (2007).

44. Chalavi, S., et al. The neurochemical basis of the contextual interference effect. Neurobiol Aging 66, 85–96 (2018).

45. Hermans, L., et al. Brain GABA Levels Are Associated with Inhibitory Control Deficits in Older Adults. J Neurosci 38, 7844–7851 (2018).

46. Simmonite, M., et al. Age-Related Declines in Occipital GABA are Associated with Reduced Fluid Processing Ability. Acad Radiol 26, 1053–1061 (2019).

47. Gao, F., et al. Edited magnetic resonance spectroscopy detects an age-related decline in brain GABA levels. Neuroimage 78, 75–82 (2013).

48. Porges, E.C., et al. Frontal Gamma-Aminobutyric Acid Concentrations Are Associated With Cognitive Performance in Older Adults. Biol Psychiatry Cogn Neurosci Neuroimaging 2, 38–44 (2017).

49. Cassady, K., et al. Sensorimotor network segregation declines with age and is linked to GABA and to sensorimotor performance. Neuroimage 186, 234–244 (2019).

50. Bailey, J.N., et al. Hypothesis-independent pathway analysis implicates GABA and acetyl-CoA metabolism in primary open-angle glaucoma and normal-pressure glaucoma. Hum Genet 133, 1319–1330 (2014).

51. Moreno, M.C., et al. Effect of ocular hypertension on retinal GABAergic activity. Neurochem Int 52, 675–682 (2008).

52. Zhou, X., et al. Differential Modulation of GABA_A_ and NMDA Receptors by an alpha7-nicotinic Acetylcholine Receptor Agonist in Chronic Glaucoma. Front Mol Neurosci 10, 422 (2017).

53. Najm, R., Jones, E.A. & Huang, Y. Apolipoprotein E4, inhibitory network dysfunction, and Alzheimer’s disease. Mol Neurodegener 14, 24 (2019).

54. Lunghi, C., Emir, U.E., Morrone, M.C. & Bridge, H. Short-term monocular deprivation alters GABA in the adult human visual cortex. Curr Biol 25, 1496–1501 (2015).

55. Loerch, P.M., et al. Evolution of the aging brain transcriptome and synaptic regulation. PLoS One 3, e3329 (2008).

56. Stanley, D.P. & Shetty, A.K. Aging in the rat hippocampus is associated with widespread reductions in the number of glutamate decarboxylase-67 positive interneurons but not interneuron degeneration. J Neurochem 89, 204–216 (2004).

57. Caspary, D.M., Milbrandt, J.C. & Helfert, R.H. Central auditory aging: GABA changes in the inferior colliculus. Exp Gerontol 30, 349–360 (1995).

58. Hua, T., Kao, C., Sun, Q., Li, X. & Zhou, Y. Decreased proportion of GABA neurons accompanies age-related degradation of neuronal function in cat striate cortex. Brain Res Bull 75, 119–125 (2008).

59. Stagg, C.J., Bachtiar, V. & Johansen-Berg, H. What are we measuring with GABA magnetic resonance spectroscopy? Commun Integr Biol 4, 573–575 (2011).

60. Hendry, S.H., Fuchs, J., deBlas, A.L. & Jones, E.G. Distribution and plasticity of immunocytochemically localized GABA_A_ receptors in adult monkey visual cortex. J Neurosci 10, 2438–2450 (1990).

61. Wu, G.K., Arbuckle, R., Liu, B.H., Tao, H.W. & Zhang, L.I. Lateral sharpening of cortical frequency tuning by approximately balanced inhibition. Neuron 58, 132–143 (2008).

62. Leventhal, A.G., Wang, Y., Pu, M., Zhou, Y. & Ma, Y. GABA and its agonists improved visual cortical function in senescent monkeys. Science 300, 812–815 (2003).

63. Cook, E., Hammett, S.T. & Larsson, J. GABA predicts visual intelligence. Neurosci Lett 632, 50–54 (2016).

64. Stagg, C.J., Bachtiar, V. & Johansen-Berg, H. The role of GABA in human motor learning. Curr Biol 21, 480–484 (2011).

65. Boroojerdi, B., Battaglia, F., Muellbacher, W. & Cohen, L.G. Mechanisms underlying rapid experience-dependent plasticity in the human visual cortex. Proc Natl Acad Sci U S A 98, 14698–14701 (2001).

66. Park, S., et al. Shorter sleep duration is associated with lower GABA levels in the anterior cingulate cortex. Sleep Med 71, 1–7 (2020).

67. Qiu, M., Ramulu, P.Y. & Boland, M.V. Association Between Sleep Parameters and Glaucoma in the United States Population: National Health and Nutrition Examination Survey. Journal of Glaucoma 28, 97–104 (2019).

68. Wang, H., Zhang, Y., Ding, J. & Wang, N. Changes in the circadian rhythm in patients with primary glaucoma. PLoS One 8, e62841 (2013).

69. Carp, J., Park, J., Polk, T.A. & Park, D.C. Age differences in neural distinctiveness revealed by multi-voxel pattern analysis. Neuroimage 56, 736–743 (2011).

70. Bowman, C.R., Chamberlain, J.D. & Dennis, N.A. Sensory Representations Supporting Memory Specificity: Age Effects on Behavioral and Neural Discriminability. J Neurosci 39, 2265–2275 (2019).

71. Atallah, B.V. & Scanziani, M. Instantaneous modulation of gamma oscillation frequency by balancing excitation with inhibition. Neuron 62, 566–577 (2009).

72. Haider, B., Duque, A., Hasenstaub, A.R. & McCormick, D.A. Neocortical network activity in vivo is generated through a dynamic balance of excitation and inhibition. J Neurosci 26, 4535–4545 (2006).

73. Dudek, F.E. & Sutula, T.P. Epileptogenesis in the dentate gyrus: a critical perspective. Prog Brain Res 163, 755–773 (2007).

74. Findley, C.A., Bartke, A., Hascup, K.N. & Hascup, E.R. Amyloid Beta-Related Alterations to Glutamate Signaling Dynamics During Alzheimer’s Disease Progression. ASN Neuro 11, 1759091419855541 (2019).

75. Siskova, Z., et al. Dendritic structural degeneration is functionally linked to cellular hyperexcitability in a mouse model of Alzheimer’s disease. Neuron 84, 1023–1033 (2014).

76. Hascup, K.N. & Hascup, E.R. Altered neurotransmission prior to cognitive decline in AbetaPP/PS1 mice, a model of Alzheimer’s disease. J Alzheimers Dis 44, 771–776 (2015).

77. Minkeviciene, R., et al. Age-related decrease in stimulated glutamate release and vesicular glutamate transporters in APP/PS1 transgenic and wild-type mice. J Neurochem 105, 584–594 (2008).

78. Rupsingh, R., Borrie, M., Smith, M., Wells, J.L. & Bartha, R. Reduced hippocampal glutamate in Alzheimer disease. Neurobiol Aging 32, 802–810 (2011).

79. Kantarci, K., et al. Proton MR spectroscopy in mild cognitive impairment and Alzheimer disease: comparison of 1.5 and 3 T. AJNR Am J Neuroradiol 24, 843–849 (2003).

80. Dreyer, E.B., Zurakowski, D., Schumer, R.A., Podos, S.M. & Lipton, S.A. Elevated glutamate levels in the vitreous body of humans and monkeys with glaucoma. Arch Ophthalmol 114, 299–305 (1996).

81. Brooks, D.E., Garcia, G.A., Dreyer, E.B., Zurakowski, D. & Franco-Bourland, R.E. Vitreous body glutamate concentration in dogs with glaucoma. Am J Vet Res 58, 864–867 (1997).

82. Carter-Dawson, L., et al. Vitreal glutamate concentration in monkeys with experimental glaucoma. Invest Ophthalmol Vis Sci 43, 2633–2637 (2002).

83. Honkanen, R.A., et al. Vitreous amino acid concentrations in patients with glaucoma undergoing vitrectomy. Arch Ophthalmol 121, 183–188 (2003).

84. Gramlich, O.W., Godwin, C.R., Wadkins, D., Elwood, B.W. & Kuehn, M.H. Early Functional Impairment in Experimental Glaucoma Is Accompanied by Disruption of the GABAergic System and Inceptive Neuroinflammation. Int J MolSci 22(2021).

85. Zhou, X., Zhang, R., Zhang, S., Wu, J. & Sun, X. Activation of 5-HT1A Receptors Promotes Retinal Ganglion Cell Function by Inhibiting the cAMP-PKA Pathway to Modulate Presynaptic GABA Release in Chronic Glaucoma. J Neurosci 39, 1484–1504 (2019).

86. Zhou, X., et al. Alpha7 nicotinic acetylcholine receptor agonist promotes retinal ganglion cell function via modulating GABAergic presynaptic activity in a chronic glaucomatous model. Sci Rep 7, 1734 (2017).

87. Provencher, S.W. Automatic quantitation of localized in vivo 1H spectra with LCModel. NMR Biomed 14, 260–264 (2001).

88. Provencher, S.W. Estimation of metabolite concentrations from localized in vivo proton NMR spectra. Magn Reson Med 30, 672–679 (1993).

89. Trivedi, V., et al. Widespread brain reorganization perturbs visuomotor coordination in early glaucoma. Sci Rep 9, 14168 (2019).

90. Brainard, D.H. The Psychophysics Toolbox. Spat Vis 10, 433–436 (1997).

